# Mitigating biases of rescaling in forward-in-time population genetic simulations

**DOI:** 10.64898/2026.09.24.754284

**Authors:** Takahiro Sakamoto

## Abstract

Forward-in-time population genetic simulations are widely used in evolutionary analyses, but simulating large populations and long genomic regions remains computationally demanding. To reduce this cost, parameter rescaling is widely employed, in which the original evolutionary process is approximated by one with a smaller population size and fewer generations. Recently, several studies using the SLiM simulator have raised concerns about the accuracy of this rescaling approach. In this study, we show that many of the biases reported in these studies can be mitigated by using a different simulation algorithm. These results reveal that the accuracy of parameter rescaling depends on how well the simulation algorithm preserves diffusion-limit properties under rescaling.

## Introduction

Evolutionary simulations are becoming an indispensable tool in population genetic studies, ranging from theoretical investigations to statistical inference based on empirical data (Adrion *et al*. 2020). Among these approaches, forward-in-time simulations are particularly useful for accommodating complex selection, population structure, and mating systems, and they have become increasingly popular with the recent development of userfriendly software packages including SLiM (Haller and Messer 2019; Haller et al. 2019) and fwdpy (Thornton 2014). However, a major drawback of forward simulations is their high computational cost, which increases rapidly with population and genome size, thereby imposing practical limitations on many applications.

To reduce this high computational cost, many studies have employed an approximation method known as parameter rescaling. Under this method, the evolution of a population with size *N*, selection coefficient *s*, mutation rate *µ*, and recombination rate *r* over *g* generations is approximated by the dynamics of a rescaled population with size *N*_rescaled_ = *N*/*Q*, selection coefficient *s*_rescaled_ = *sQ*, mutation rate *µ*_rescaled_ = *µQ*, recombination rate *r*_rescaled_ = *rQ*, evolving for *g*_rescaled_ = *g*/*Q* generations, where *Q* denotes the rescaling factor. This transformation allows the process to be simulated with smaller population sizes and fewer generations, thereby greatly reducing the computational burden. This strategy is motivated by the fact that, under diffusion-limit approximations (i.e., continuous time and continuous allele-frequency limits), evolutionary dynamics depend only on population-size-scaled parameters (Hill and Robertson 1966; Ewens 2004). Although such rescaling is not exact in the discrete-generation framework of forward simulations, it is generally considered a valid approximation when rescaled parameters remain sufficiently small for the diffusion approximation to hold (Johri et al. 2026).

However, several recent studies have shown that rescaling can significantly bias evolutionary dynamics (Dabi and Schrider 2025; Ferrari et al. 2025; Marsh et al. 2026) (see also Uricchio and Hernandez 2014). Using the SLiM simulator, these studies reported substantial discrepancies between rescaled and unscaled simulations across a range of evolutionary scenarios. Deviations are observed in many summary statistics including nucleotide diversity, site-frequency spectra (SFS), strength of linkage disequilibrium (LD), and fixation time. Marsh *et al*. (2026) suggested that rescaling can lead to biases when recombination rates are high or genomic regions are sufficiently long so that multiple crossovers are likely to occur within a simulated region (see also Johri *et al*. 2026). Marsh *et al*. (2026) also pointed out that some of the previously observed discrepancies arise from the excessively large selection coefficients after the rescaling is applied, which violates the assumptions underlying the diffusion approximation.

Building on Marsh *et al*. (2026) and Johri *et al*. (2026), we consider two main causes of the inaccuracy of rescaled simulations under appropriate selection strength. First, long-range LD is overrepresented in rescaled SLiM simulations because the amount of LD that can be resolved within a single generation is limited. To illustrate this, we consider a rescaled simulation witha rescaling factor *Q* = 4 (Figure 1a). Under this setting, time is compressed such that one generation in the rescaled simulation corresponds to four generations in the original dynamics, while the recombination rate is multiplied by four to preserve the expected number of recombination events. Under the original timescale, haplotype pairs are reshuffled in each generation, and recombination events occur between different haplotypes over time (top row, Figure 1a). As a result, haplotypes after four generations become mosaics of genomic regions derived from multiple parents in the initial state. In contrast, in rescaled SLiM simulations, all recombination events within a single “compressed” generation occur between the same pair of haplotypes carried by a diploid individual. This prevents each haplotype from inheriting genomic regions from more than two parental haplotypes, thereby limiting the resolution of long-range LD (middle row, Figure 1a). This increased linkage can lead to reduced genetic diversity through stronger linked selection and increased interference among selected variants.

**Figure 1.**
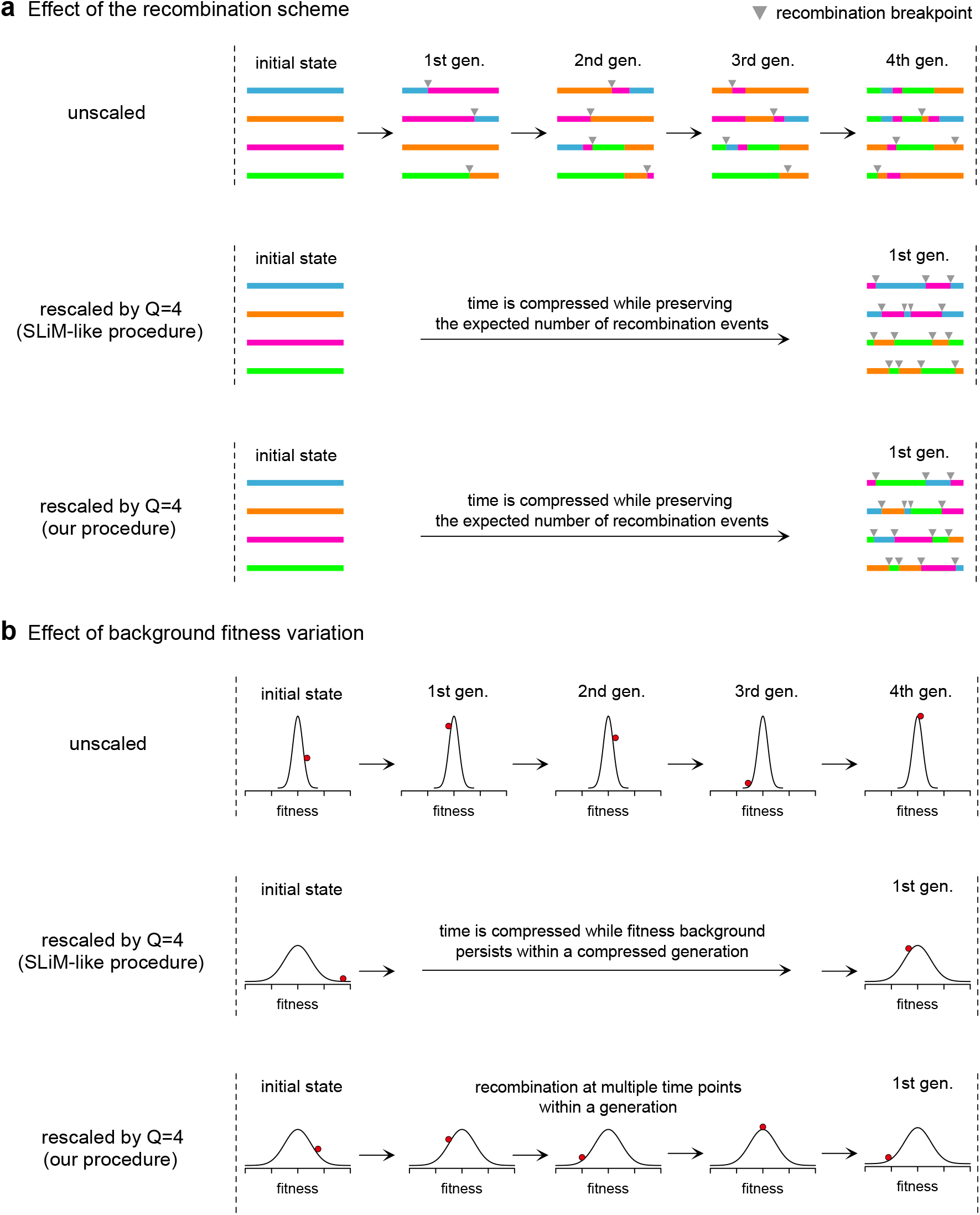
Potential biases in rescaled simulations and our proposed solution. (a) Effect of the recombination scheme. In this example, a rescaling factor of *Q* = 4 is assumed. The top row illustrates how recombination shuffles genomes over *Q* generations in the original timescale. By generation *Q*, each haplotype is composed of segments with mixed origins across the entire region. The second row shows a schematic view of the rescaled simulations using a SLiM-like recombination procedure. Here, one generation in the rescaled simulation corresponds to *Q* generations in the original timescale. Although the expected number of recombination events is preserved by multiplying the recombination rate by *Q*, all recombination events contributing to a given offspring occur between the same pair of parental haplotypes, failing to reproduce the well-shuffled ancestry observed in the original timescale. The bottom row illustrates the present method. By selecting recombining haplotypes independently at each recombination event, this approach allows mixing of ancestry across the genome while minimally affecting the selection strength and the recombination rate. (b) Effect of the background fitness variation. In the original timescale, the genetic background changes every generation, and its effects are averaged over time. In contrast, in a SLiM-like rescaled simulation, the same background persists for *Q* generations in the original timescale, allowing variants in high-fitness backgrounds to produce disproportionate numbers of offspring, especially for large *Q* (Marsh et al. 2026). This increases stochasticity in allele frequency dynamics and reduces the effective population size. In our method, multiple recombination events within a generation reshuffle genetic backgrounds and prevent such persistence.

Second, in the presence of many unlinked selected variants,rescaling can skew the offspring distribution, resulting in increased stochasticity (Marsh et al. 2026). This effect is illustrated in Figure 1b by focusing on the fate of an allele indicated by a red circle. Because many selected variants are unlinked, the fitness background of this allele varies from generation to generation. Its effects are averaged over time in the original-scale simulation, therefore it does not contribute to the variance of offspring number among alleles very much in the long run. In contrast, in rescaled simulations, the same fitness background is fixed within a “compressed” generation. In other words, the variant in a high-fitness background can enjoy this advantage for a period corresponding to *Q* generations on the original timescale, which may result in bursts of offspring production. Thus, in addition to the random genetic drift due to finite population size, variation in fitness background introduces an additional source of stochasticity, leading to reduced effective population sizes and lower genetic diversity.

While these explanations are reasonable, it remains unclear whether these two factors sufficiently explain the biases observed in rescaled SLiM simulations. It also remains unexplored whether it is possible to mitigate these biases by using a simulation procedure different from that implemented in SLiM.

In this study, we present a new simulator that mitigates biases introduced by parameter rescaling in forward-in-time simulations. Rather than aiming to reproduce biological processes in detail like SLiM, this approach is designed to preserve the assumptions underlying diffusion-limit approximations under rescaling. The two main sources of bias are addressed by modifying the simulation scheme in ways that do not strictly adhere to biological realism. Using this framework, we show that these modifications substantially reduce rescaling-induced biases across a wide range of rescaling factors and genomic conditions.

## Methods

### Model

We consider 2*N* haploid individuals, or equivalently *N* diploid individuals under codominance. The population size is constant, and time is measured in generations in the scale of the Wright–Fisher model, denoted by *t*. We assume additive log fitness across loci, such that an individual’s log fitness is the sum of the effects of all variants it carries. The simulated region has length *L*. The recombination and mutation rates per site per generation are denoted by *r* and *µ*, respectively.

### New simulation algorithm

In this simulator, we aim to mitigate rescaling-induced biases by introducing (i) recombination among more than two haplotypes in a generation (bottom row; Figure 1a) and (ii) multiple recombination events within a generation to reshuffle fitness backgrounds in a finer time resolution (bottom row; Figure 1b). Although these mechanisms do not strictly follow the biological meiotic process, they directly address the sources of bias described above and may lead to more accurate rescaled simulations. To accommodate these extensions, we formulate the model based on a Moran framework rather than the Wright–Fisher model. Note that parameters are still defined according to the Wright–Fisher model.

The simulator consists of two types of events: replacement and recombination-mutation events. Each event is constructed to result in the same diffusion limit as the standard Wright–Fisher model.

#### Replacement event

In a replacement event, one individual is replaced by a new offspring, thereby implementing selection and random genetic drift. Let *w*_*i*_ denote the log fitness of individual *i*. An individual to be removed is chosen with probability proportional to 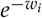,

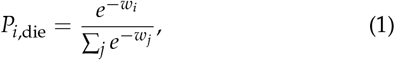

and an individual to reproduce is chosen with probability proportional to 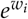,

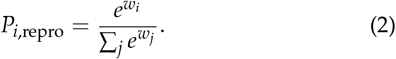

The genome of the reproducing individual is copied and replaces that of the removed individual.

To quantify allele frequency changes through this event, we consider an allele with selection coefficient *s*. Letting *p* be its allele frequency, mean and variance of the frequency change is given by

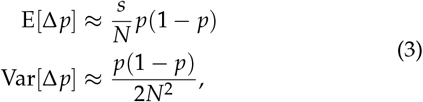

assuming weak selection and linkage equilibrium (Appendix A).

Equation 3 implies that a single replacement event introduces the same amount of stochasticity as 1/*N* generations in the Wright–Fisher model. The expected allele frequency change per Wright–Fisher generation is *sp*(1 − *p*), consistent with the standard Wright–Fisher model.

#### Recombination-mutation event

This event first reshuffles haplotypes through recombination and then introduces new mutations. The infinite-sites model (Kimura 1969) is assumed, such that mutation positions and recombination breakpoints are continuous and take values in (0, *L*). In an event, we introduce recombinations and mutations corresponding to Δ*t* Wright–Fisher generations.

First, the total number of recombinations in the population is drawn from a Poisson distribution with mean *NrL*Δ*t*. For each recombination, two haplotypes are chosen at random from all haplotypes and recombined at a breakpoint drawn from Uniform(0, *L*). A haplotype can engage in multiple recombination events with different haplotypes in a generation. Since each recombination involves two haplotypes, the expected recombination rate per haplotype per generation is *rL*, consistent with the standard Wright–Fisher model.

Next, new mutations are introduced. The total number of mutations in the population is drawn from a Poisson distribution with mean 2*NµL*Δ*t*, and each mutation is assigned to a randomly chosen haplotype. This gives the same expected number of new mutations as the standard Wright–Fisher model.

#### Frequency of recombination-mutation event

An important modeling choice involves how frequently recombination-mutation event should occur. Although a natural implementation would apply a mutation-recombination event with Δ*t* = 1/*N* after each replacement event, mutation-recombination events are computationally intensive, as they involve copying mutations on the recombining haplotypes. Thus, it is preferable to reduce their frequency as long as this does not introduce biases due to persistent fitness backgrounds. Practically, it can be implemented by running a recombination-mutation event after several replacement events.

To quantify how large Δ*t* is allowed without introducing biases, we approximate the additional stochasticity introduced by fixed fitness backgrounds as a function of Δ*t* (Appendix B). The additional variance per generation is approximated by

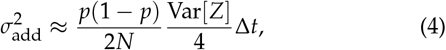

where 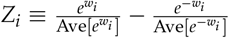 and Var[*Z*] is its variance. When |*w*_*i*_ |*≪* 1 and Δ*t* = 1, Equation 4 reduces to the additional variance expected in the Wright–Fisher model in the presence of fitness variation (Charlesworth 2012), by noting *Z*_*i*_ *≈* 2*w*_*i*_.

This equation shows that the deviation arising from fixed fitness background can be mitigated by assuming small Δ*t*, consistent with our intuition (Figure 1b). As the amount of random genetic drift in the Wright–Fisher diffusion process is 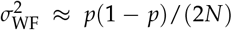 per generation, for *σ*^2^ to be negligible compared with this, we require Var[*Z*]Δ*t*/4 *≪* 1. In this paper, we determine Δ*t ≈* min(0.001 *×* 4/Var[*Z*], 1) such that 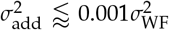. In simulations, we calculate Var[*Z*] every Wright–Fisher generation and determine the interval of recombination-mutation event, Δ*t*, following this method.

### Simulation settings

To examine the extent to which our implementation mitigates rescaling-induced biases, we performed extensive simulations. We considered two scenarios: (I) purifying selection with constant selective effects and (II) purifying selection together with positive selection. For ease of comparison, parameter values were chosen to match those in Marsh *et al*. (2026). We assumed a population size of 2*N* = 2 *×*10^6^, a mutation rate of *µ* = 3 *×*10^−9^ per site per generation, and a recombination rate of *r* = 1*×*10^−8^ per site per generation.

In the purifying-selection scenario (scenario I), a new mutation was neutral with probability 0.5 and deleterious otherwise, with selection coefficient 4*Ns* = −100. Because *s* is defined here as the heterozygous effect, whereas it is defined as the homozygous effect in Marsh *et al*. (2026), this setting corresponds to the strong purifying-selection scenario in Marsh *et al*. (2026).

In the scenario combining purifying selection and positive selection (scenario II), a new mutation was neutral with probability 0.5, deleterious with probability 0.4999, and advantageous with probability 0.0001. All deleterious mutations had the same selection coefficient, 4*Ns* = −100, and the selection coefficient of a beneficial mutation was drawn from an exponential distribution with mean 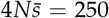. This setting corresponds to the recurrent selective-sweep scenario in Marsh *et al*. (2026) ^1^.

We considered six lengths of genomic regions: *L* = 10 kb, 40 kb, 200 kb, 1 Mb, 5 Mb, and 25 Mb. We also considered five rescaling factors, *Q* = 25, 50, 125, 250, and 625. However, because small rescaling factors combined with long genomic regions impose prohibitive computational costs, we examined only a subset of the combinations of *L* and *Q*. The examined pairs are summarized in Table S1 and are identical to those examined in Marsh *et al*. (2026).

In addition to our custom-code simulation, we also ran models with these parameter settings using SLiM v5.2 (Haller *et al*. 2026). These simulations are used to clarify how the implementation difference affects the outcome.

### Statistics

To summarize the simulation dynamics, we focused on several summary statistics: nucleotide diversity (*π*; Nei and Li (1979)), site-frequency spectrum (SFS; Watterson (1975)), squared linkage disequilibrium (*r*^2^; Hill and Robertson (1968)), the number of fixed mutations, and fixation time. In this study, the SFS was normalized so that its sum was one. We ran each simulation for 14*N*_rescaled_ generations and sampled 100 haplotypes without replacement, corresponding to 50 diploid individuals, from which *π*, SFS, and LD were calculated. For LD, average *r*^2^ was calculated for non-overlapping bins with a width of 250 bp. When the number of segregating variants exceeded 100,000, 100,000 variants were randomly subsampled for LD calculations. We also recorded all mutations that are fixed during the last 4*N*_rescaled_, from which the number of fixed mutations and fixation time were calculated.

For each combination of *L* and *Q*, we ran 50 simulation replicates to evaluate mean value as well as standard deviation across simulation replicates.

## Results

### Purifying selection scenario

We first examine cases in which all selected mutations are deleterious (scenario I). In general, we observed notable effects of the rescaling factor for long genomic segments (≥ 1 Mb) in the SLiM simulations whereas these biases are largely mitigated in our implementation.

As an example, Figure 2 shows nucleotide diversity calculated from neutral alleles. Compared with theoretical prediction calculated based on Nordborg *et al*. (1996) (horizontal dashed lines, see Appendix C for details), neutral nucleotide diversity is substantially reduced for *Q* = 250 or 625 in SLiM simulations with long segments (i.e., ≥1 Mb), quantitatively consistent with Marsh *et al*. (2026). In contrast, our simulator retains an amount of variation close to the theoretical expectation, although a small decrease is observed for *Q* = 625. For example, in a 25 Mb segment with a rescaling factor *Q* = 625, SLiM simulation retains only 9.1 % of variation predicted by theory, whereas our simulator maintains 93.1 % of variation.

**Figure 2.**
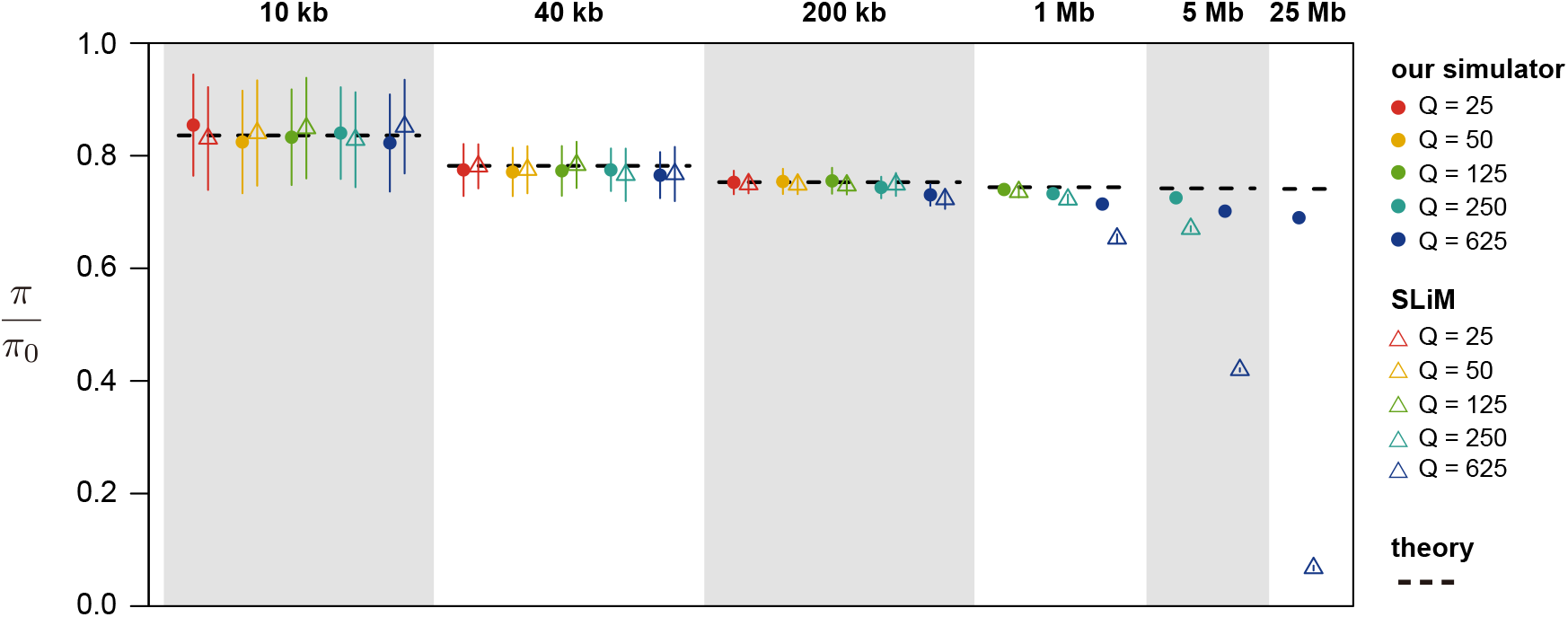
Nucleotide diversity under purifying selection. Nucleotide diversity of neutral variants, normalized by the neutral expectation without linked selection (*π*_0_ = 4*NµL ×*0.5), is shown for different genomic region lengths and rescaling factors *Q*. Filled circles and open triangles represent results from our simulator and SLiM, respectively. Points and error bars indicate the mean and standard deviation across 50 simulation replicates. Dashed lines show the theoretical expectation based on Nordborg *et al*. (1996).

In Supplementary Figures, we also compare other statistics and find notable biases in the SLiM simulations in the parameter region identified above. Overall, for a long genomic region with a large rescaling factor, SLiM shows an increased proportion of neutral singletons (Figure S1b), a decreased proportion of rare deleterious mutations (Figure S2b), elevated linkage disequilibrium (Figure S3b), elevated fixation probability of deleterious mutations (Figure S4b), and shorter fixation times of neutral mutations (Figure S5b). These patterns are largely consistent with the increased stochasticity (i.e., reduced effective population size). By contrast, these biases are largely mitigated in our implementation (Figures S1-S5). These results show that the robustness against the rescaling significantly depends on the implementation in simulators.

#### Scenario with both purifying and positive selection

We next introduce a small amount of beneficial mutations in addition to deleterious mutations (scenario II). Due to recurrent selective sweep, nucleotide diversity calculated from neutral variants in this scenario is smaller than scenario I (Figure S6). We again observe the reduction of diversity in SLiM simulations when assuming a long segment (*≥* 5 Mb) and a large rescaling factor (*Q* = 625). In contrast, our implementation shows largely consistent results across different rescaling factors, suggesting the biases are largely mitigated.

Other statistics show similar qualitative patterns as scenario I (Figures S7-S11). Notably, all biases are substantially reduced in our simulator. These results show that our simulation method is more robust against biases arising from parameter rescaling.

## Discussion

The present findings show that the accuracy of rescaling in forward-in-time simulations depends on the simulation implementation. Although the effect of rescaling factors can be drastic in SLiM simulations, our simulator produces largely consistent results across rescaling factors. By construction, the present algorithm addresses the two sources of potential biases in the rescaled simulations (Figure 1): insufficient resolution of long-range linkage disequilibrium and excess stochasticity caused by persistent fitness backgrounds. The good performance of our simulator suggests that these two mechanisms are major sources of the previously observed biases in the rescaled simulations.

The present simulator aims to more accurately preserve the dynamics expected under the diffusion limit even under parameter rescaling. Consequently, our implementation of recombination and mutation is not intended to reproduce the details of biological processes. This contrasts with the SLiM simulator, which prioritizes biological realism. Our results show that, when the goal is to approximate diffusion-limit dynamics under parameter rescaling, biologically realistic simulators are not necessarily the best choice.

Regarding the computational cost, we conducted a rough comparison of computational time (Appendix D). The present simulator is slower than SLiM by orders of magnitude when genomic regions are long and the rescaling factor is large, reflecting the need for frequent recombination-mutation events to maintain accuracy in our simulator. This suggests that our algorithm mitigates rescaling-induced biases at the expense of computational speed. Nevertheless, our rescaled simulations consistently become faster as the rescaling factor increases, highlighting the computational benefit of rescaling. In the future, the present algorithm and simulation code may be further refined to improve computational efficiency.

## Data Avalilability

Simulation code developed in this study is available at https://github.com/TSakamoto-evo/rescaling_simulation.

## Appendix A: Allele frequency change in a replacement event under linkage equilibrium

In this section, we derive mean and variance of allele frequency change through a replacement event under linkage equilibrium.We focus on a locus, where ancestral and derived allele have log fitness of 0 and *s*, respectively. Let *p* be the frequency of the derived allele. Under linkage equilibrium, the distribution of background log fitness is independent of the focal allele. We denote this distribution by *f* (*x*).

The probability that a derived allele dies is given by

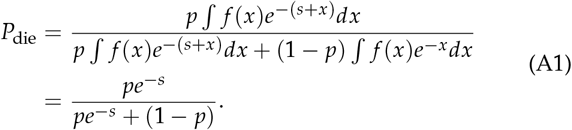

Similarly, the probability that a derived allele reproduces is

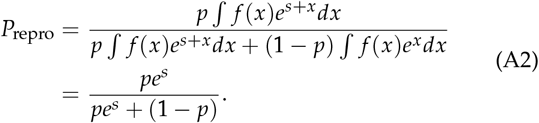

The allele frequency change satisfies

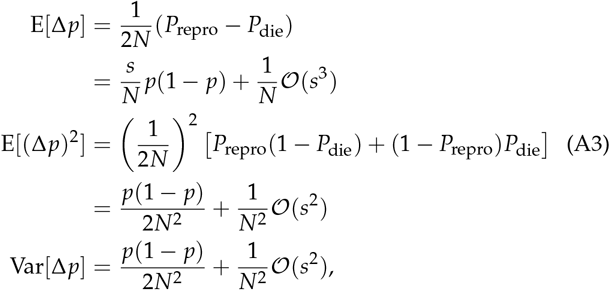

resulting in Equation 3.

## Appendix B: Stochasticity in allele frequency due to variation in background fitness

In this section, we calculate how background fitness variation affects the stochastic allele frequency change. For simplicity,we focus on the dynamics at a single neutral locus, but fitness variation exists even among the same allele because of genetic variation at other loci. Background fitness and frequency of *i*th haplotype are denoted by *w*_*i*_ and *x*_*i*_, respectively. We assume that, initially, each haplotype is present as a single copy, so that *x*_*i*_ = 1/(2*N*). Conditional on the background fitness *w*_*i*_,the expected frequency change of the *i*th haplotype in a single replacement event is approximately

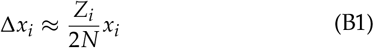

where 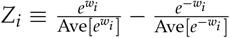. After Δ*t* Wright–Fisher generations, which is equivalent to *N*Δ*t* replacement events, the total frequency change is approximated by Δ*x*_*i*_ *≈ Z*_*i*_*x*_*i*_ Δ*t*/2.

Noting *x*_*i*_ = 1/(2*N*), the total stochastic change for *n* haplotypes sharing the same allele at the focal locus is

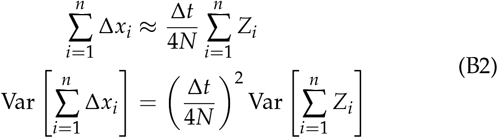

Since the *Z*_*i*_ values are defined relative to the population mean, they satisfy

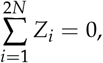

which implies

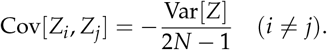

Therefore,

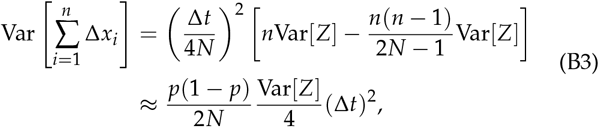

where *p* = *n*/(2*N*).

Note that the above equation quantifies the change over Δ*t* Wright–Fisher generations. By considering the stochastic change per a Wright–Fisher generation, Equation 4 is derived.

## Appendix C Theoretical prediction on the effect of background selection

As a benchmark for the nucleotide diversity observed in simulations, we calculated the theoretical expectation based on Nordborg *et al*. (1996). Equation 4 of Nordborg *et al*. (1996) approximates the reduction in nucleotide diversity due to background selection as

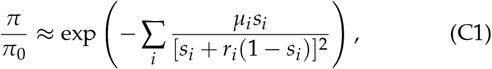

where *π*_0_ denotes the nucleotide diversity in the corresponding purely neutral model, *s*_*i*_ is the heterozygous disadvantage of deleterious mutations at the *i*th selected locus, *µ*_*i*_ is the deleterious mutation rate at that locus, and *r*_*i*_ is the recombination rate between the focal neutral site and the *i*th selected locus.

Applying this approximation to the situation considered in this study, the reduction in nucleotide diversity at position *x*(0 *≤ x ≤ L*) is approximated by

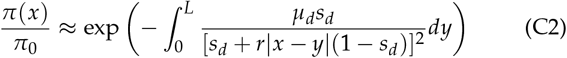

where *µ*_*d*_ = 0.5*µ* = 1.5 *×*10^−9^ is the per-site deleterious mutation rate and *s*_*d*_ = 100/(4*N*) is the heterozygous disadvantage. Averaged over the position *x*, the expected reduction is calcu-lated as

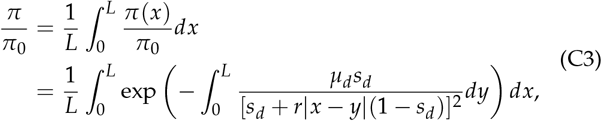

which is plotted in Figure 2.

## Appendix D: Comparison of computational time

In this section, we compare the computational time of our simulator and SLiM across different settings. As our simulator allows multiple recombination-mutation events in a generation, it is expected to slow down simulation. This is particularly the case for long genomic regions, stronger selection, and large rescaling factors, because greater variation in genetic background fitness requires more frequent reshuffling of genetic backgrounds.

Note that our purpose is not to provide a rigorous benchmark of computational efficiency between the two simulators. In addition to the model differences, it is important to acknowledge that SLiM code has been highly optimized during its development whereas our code may not be so efficient. Also, our code is specialized in the present setting whereas SLiM allows a wide variety of extensions, which may affect the difference of computational speed. Thus, the difference in computational time may not purely reflect the performance difference arising from the algorithms.

To obtain the rough computational time, we ran 10 replicates of each simulator while changing the length of a genomic region and a rescaling factor (see Table S1). We assumed the purifying selection scenario (scenario I). All replicates were run at the computer cluster SHIROKANE at the Human Genome Center (the Univ. of Tokyo) using a single CPU core of Shirokane 8 (mjob). Elapsed time (seconds) was recorded every 100 generations, and all replicates running longer than two days were stopped.

Figure S12 shows the median elapsed time across 10 replicates. In general, computational time decreases as the rescaling factor increases, consistent with the purpose of parameter rescaling. The difference between our simulator and SLiM becomes particularly pronounced when a long genomic region and a large rescaling factor are assumed. Under these parameter settings, our simulator requires more computational time because more frequent recombination-mutation events are needed to sufficiently reshuffle genetic backgrounds and prevent rescaling-induced biases. These results indicate that the increased robustness of our simulator comes at additional computational cost, partially offsetting the computational benefit of rescaling. Nevertheless, across the parameter settings examined here, simulations with larger rescaling factors are consistently faster, indicating that the computational benefit of rescaling is retained.

**Table S1.** Examined rescaling factor.

| length of genomic region ($L$) | rescaling factor ($Q$) |
| --- | --- |
| 10 kb | 25, 50, 125, 250, 625 |
| 40 kb | 25, 50, 125, 250, 625 |
| 200 kb | 25, 50, 125, 250, 625 |
| 1 Mb | 125, 250, 625 |
| 5 Mb | 250, 625 |
| 25 Mb | 625 |

**Figure S1.**
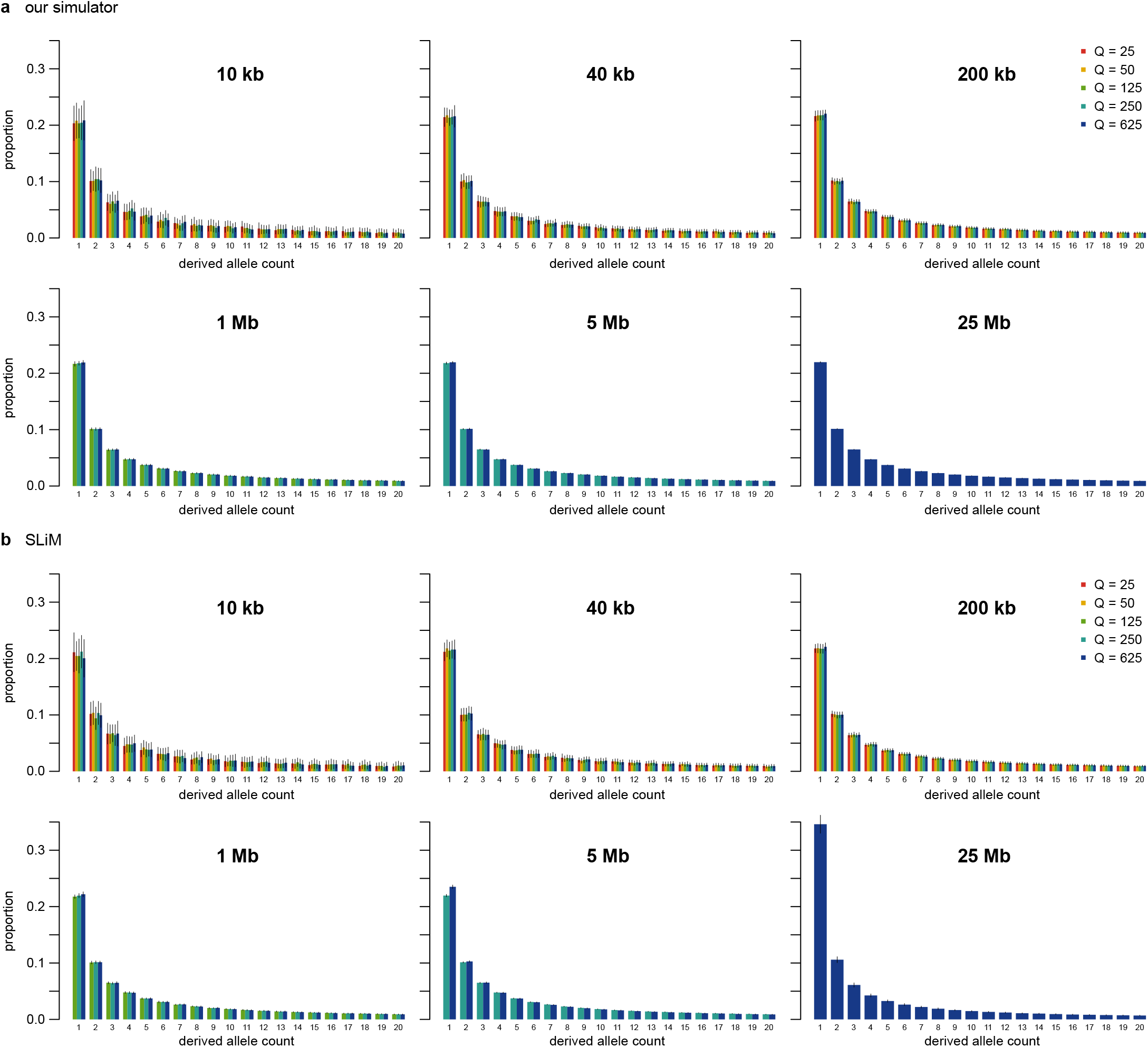
Site-frequency spectrum of neutral variants under purifying selection. The normalized site-frequency spectrum of neutral variants is shown for different genomic region lengths and rescaling factors *Q*. The spectrum was normalized within each simulation replicate so that the sum across all derived allele count classes was one. Bars and error bars indicate the mean and standard deviation across 50 simulation replicates. Panels (a) and (b) show results from our simulator and SLiM, respectively.

**Figure S2.**
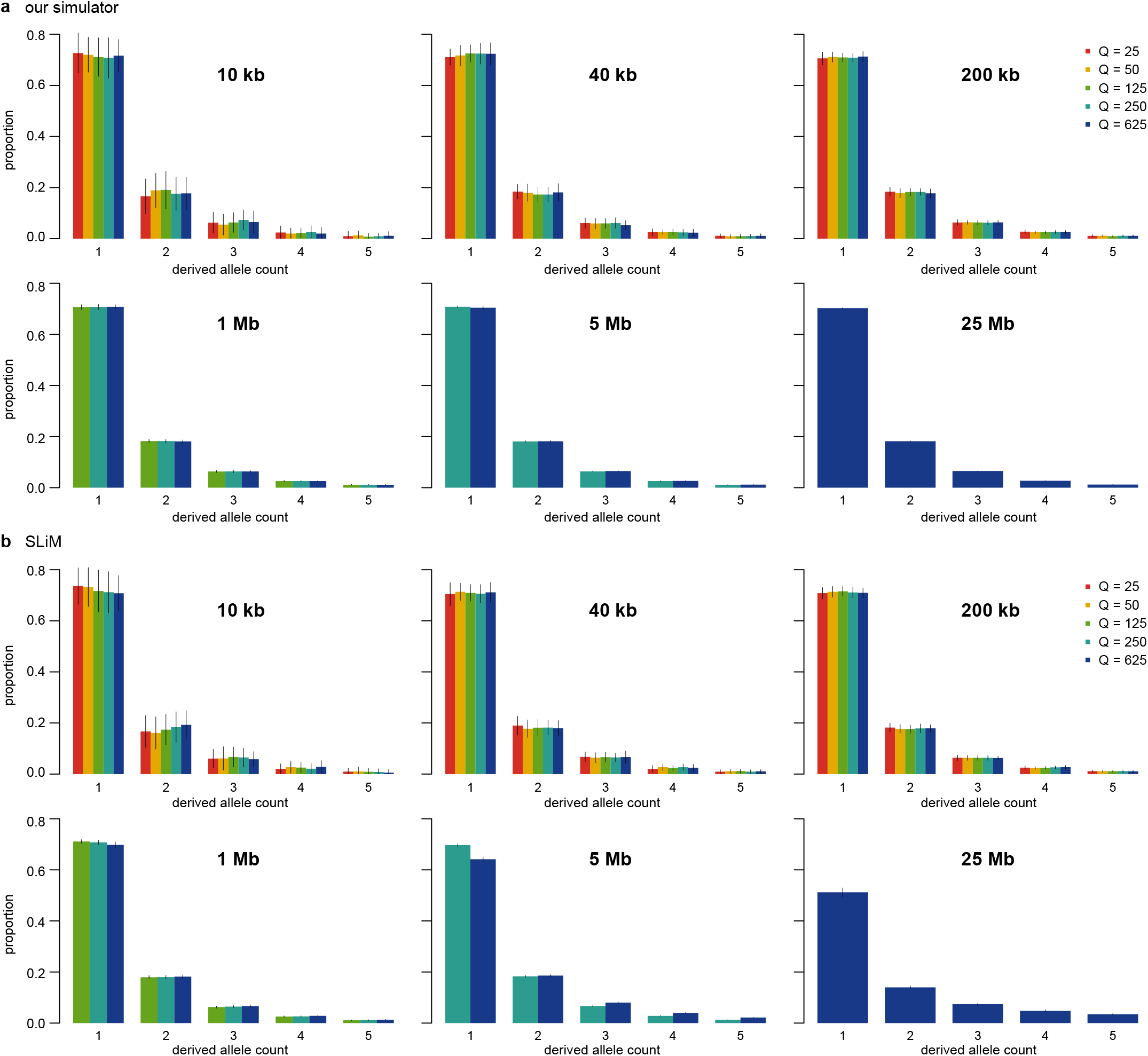
Site-frequency spectrum of deleterious variants under purifying selection. The normalized site-frequency spectrum of deleterious variants is shown for different genomic region lengths and rescaling factors *Q*. The spectrum was normalized within each simulation replicate so that the sum across all derived allele count classes was one. Bars and error bars indicate the mean and standard deviation across 50 simulation replicates. Panels (a) and (b) show results from our simulator and SLiM, respectively.

**Figure S3.**
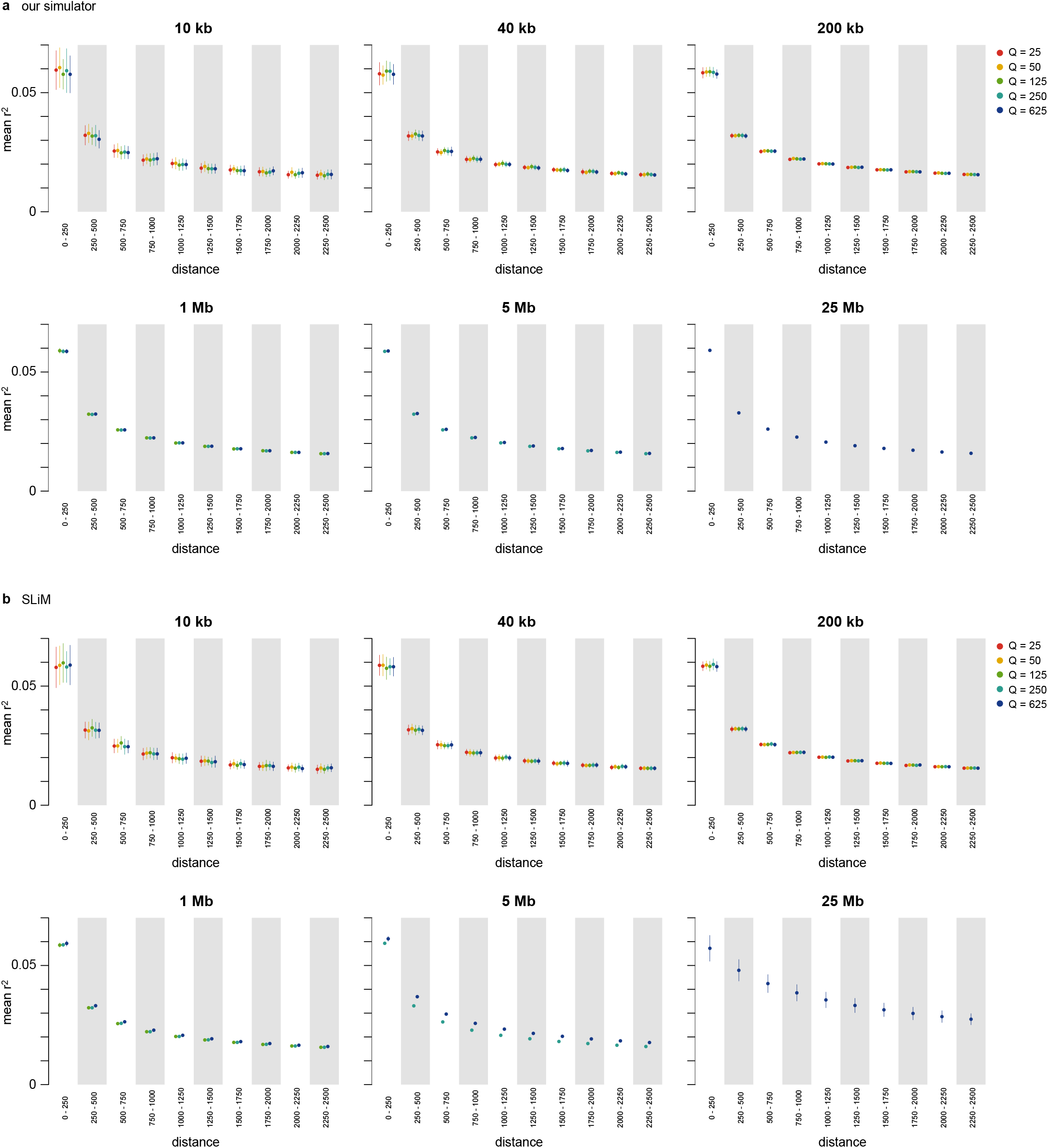
Linkage disequilibrium under purifying selection. Mean squared linkage disequilibrium (*r*^2^) is shown for different genomic region lengths and rescaling factors *Q*. Values were calculated in non-overlapping 250-bp bins. Points and error bars indicate the mean and standard deviation across 50 simulation replicates. Panels (a) and (b) show results from our simulator and SLiM, respectively.

**Figure S4.**
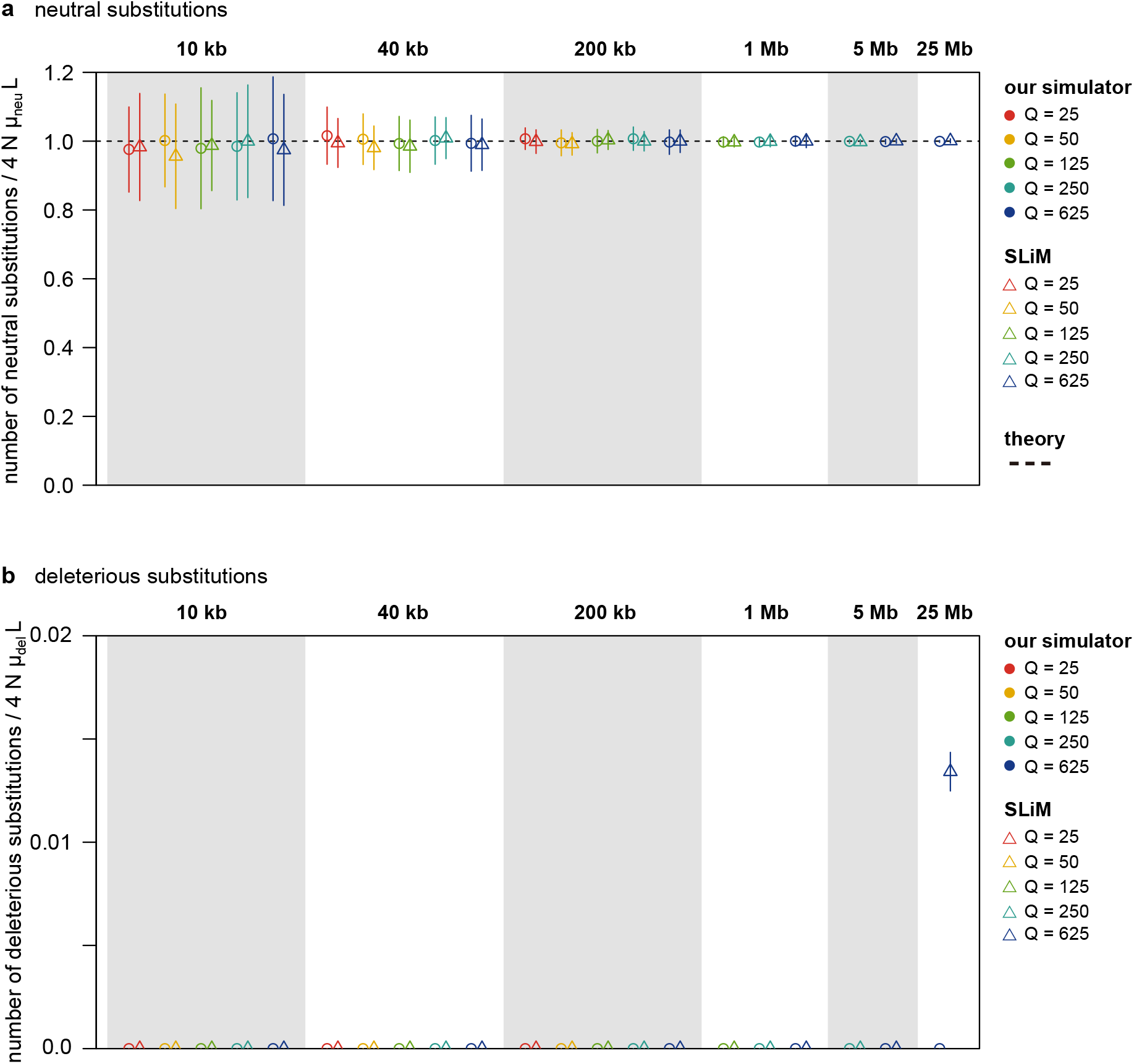
Substitution counts under purifying selection. The numbers of neutral and deleterious substitutions fixed during the last 4*N*_rescaled_ generations are shown for different genomic region lengths and rescaling factors *Q*. Counts are normalized by the corresponding neutral expectations, 4*Nµ*_neu_ *L* for neutral substitutions and 4*Nµ*_del_ *L* for deleterious substitutions. Points and error bars indicate the mean and standard deviation across 50 simulation replicates. Panel (a) shows neutral substitutions and panel (b) deleterious substitutions; in each panel, circles and triangles represent our simulator and SLiM, respectively.

**Figure S5.**
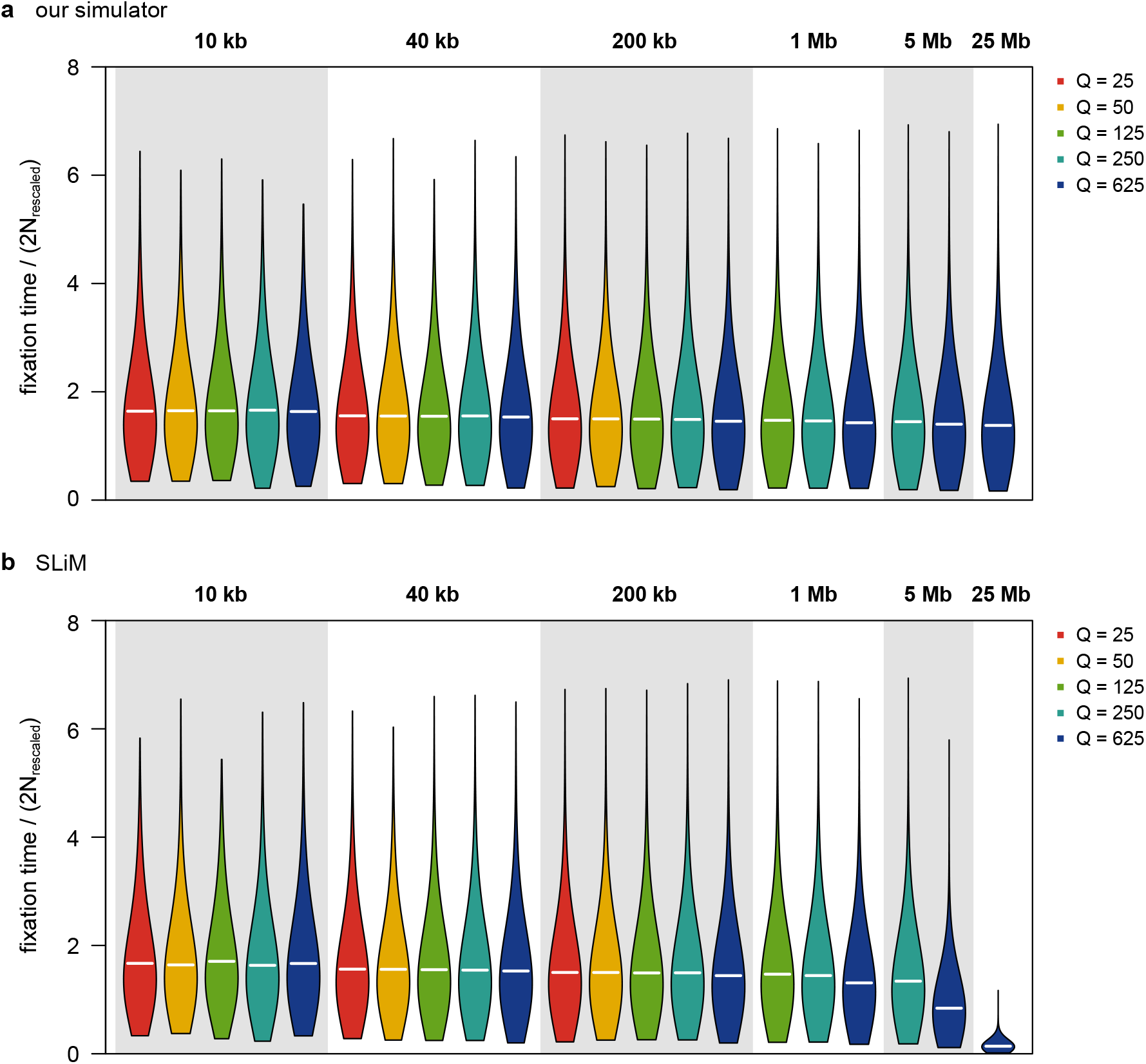
Fixation time of neutral substitutions under purifying selection. Distributions of fixation times for neutral mutations fixed during the last 4*N*_rescaled_ generations are shown for different genomic region lengths and rescaling factors *Q*. Fixation times are normalized by 2*N*_rescaled_. Horizontal white lines indicate the mean fixation time. Panels (a) and (b) show results from our simulator and SLiM, respectively.

**Figure S6.**
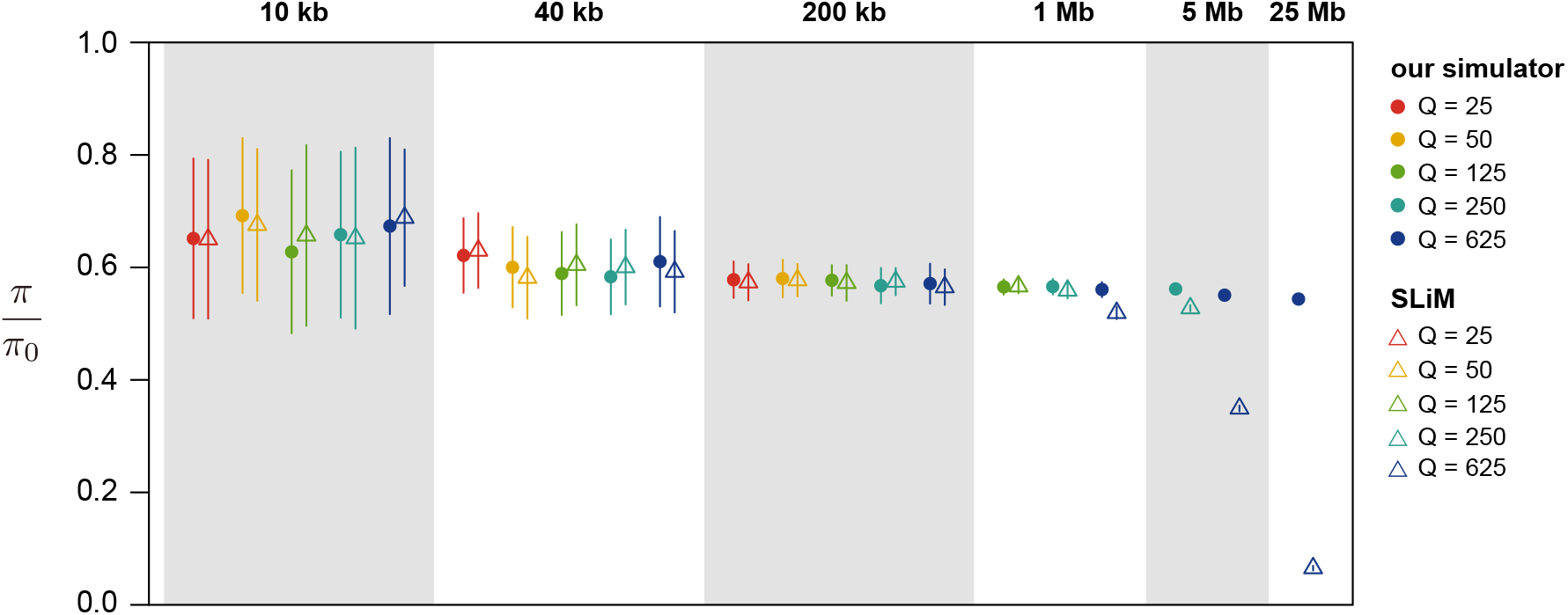
Nucleotide diversity under purifying and positive selection. Nucleotide diversity of neutral variants, normalized by the neutral expectation without linked selection (*π*_0_ = 4*NµL ×* 0.5), is shown for different genomic region lengths and rescaling factors *Q*. Filled circles and open triangles represent results from our simulator and SLiM, respectively. Points and error bars indicate the mean and standard deviation across 50 simulation replicates.

**Figure S7.**
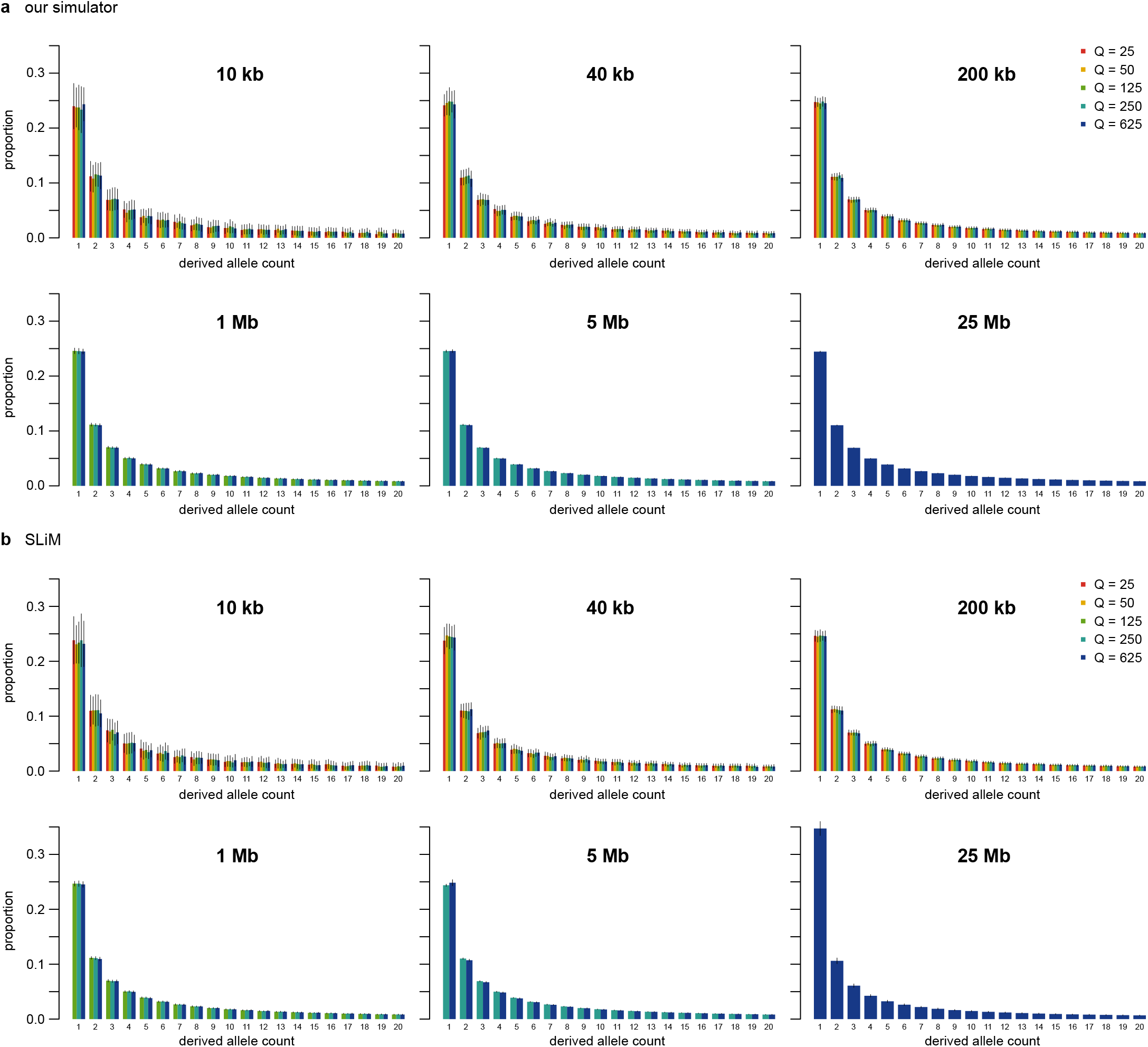
Site-frequency spectrum of neutral variants under purifying and positive selection. The normalized site-frequency spectrum of neutral variants is shown for different genomic region lengths and rescaling factors *Q*. The spectrum was normalized within each simulation replicate so that the sum across all derived allele count classes was one. Bars and error bars indicate the mean and standard deviation across 50 simulation replicates. Panels (a) and (b) show results from our simulator and SLiM, respectively.

**Figure S8.**
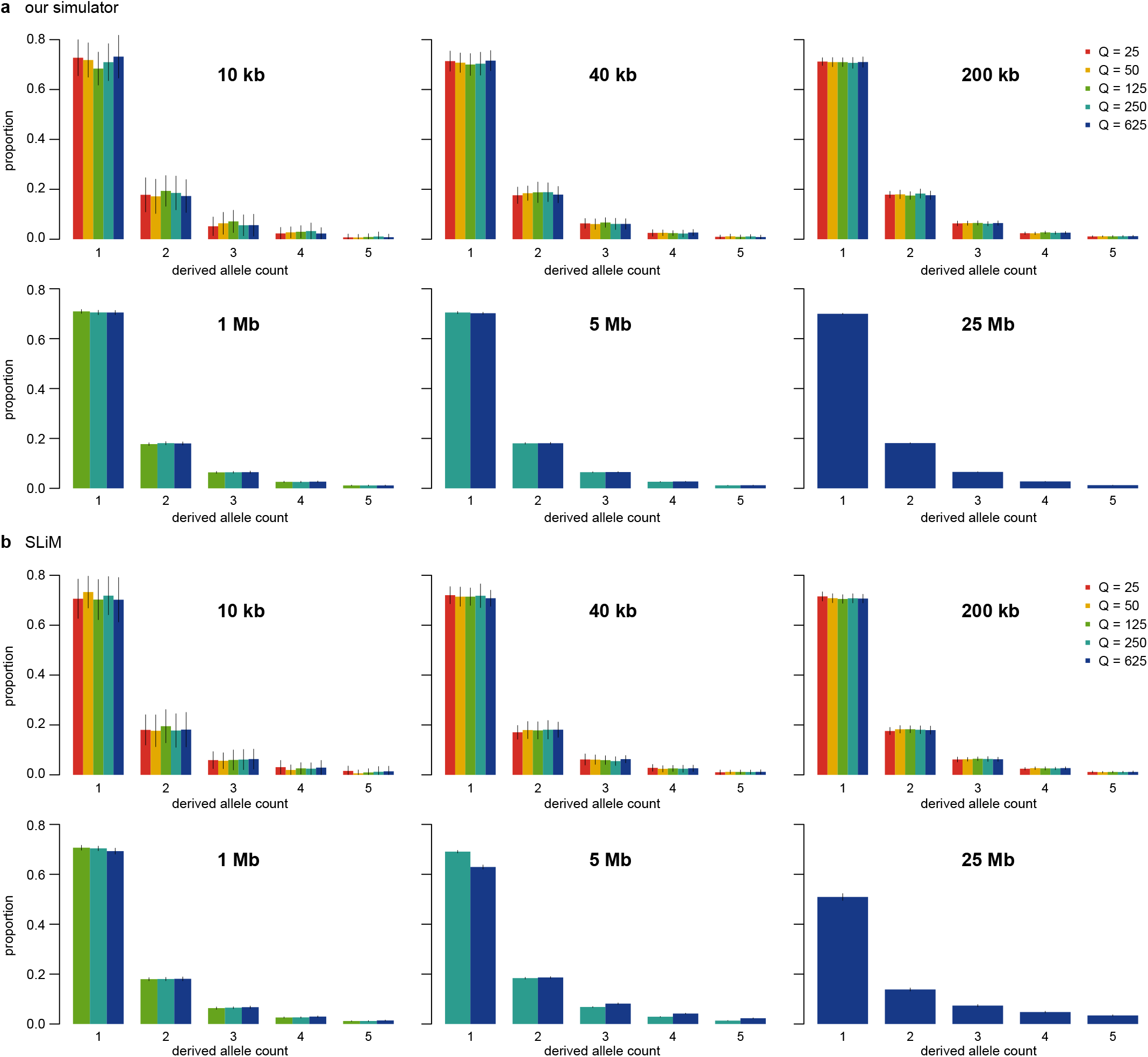
Site-frequency spectrum of deleterious variants under purifying and positive selection. The normalized site-frequency spectrum of deleterious variants is shown for different genomic region lengths and rescaling factors *Q*. The spectrum was normalized within each simulation replicate so that the sum across all derived allele count classes was one. Bars and error bars indicate the mean and standard deviation across 50 simulation replicates. Panels (a) and (b) show results from our simulator and SLiM, respectively.

**Figure S9.**
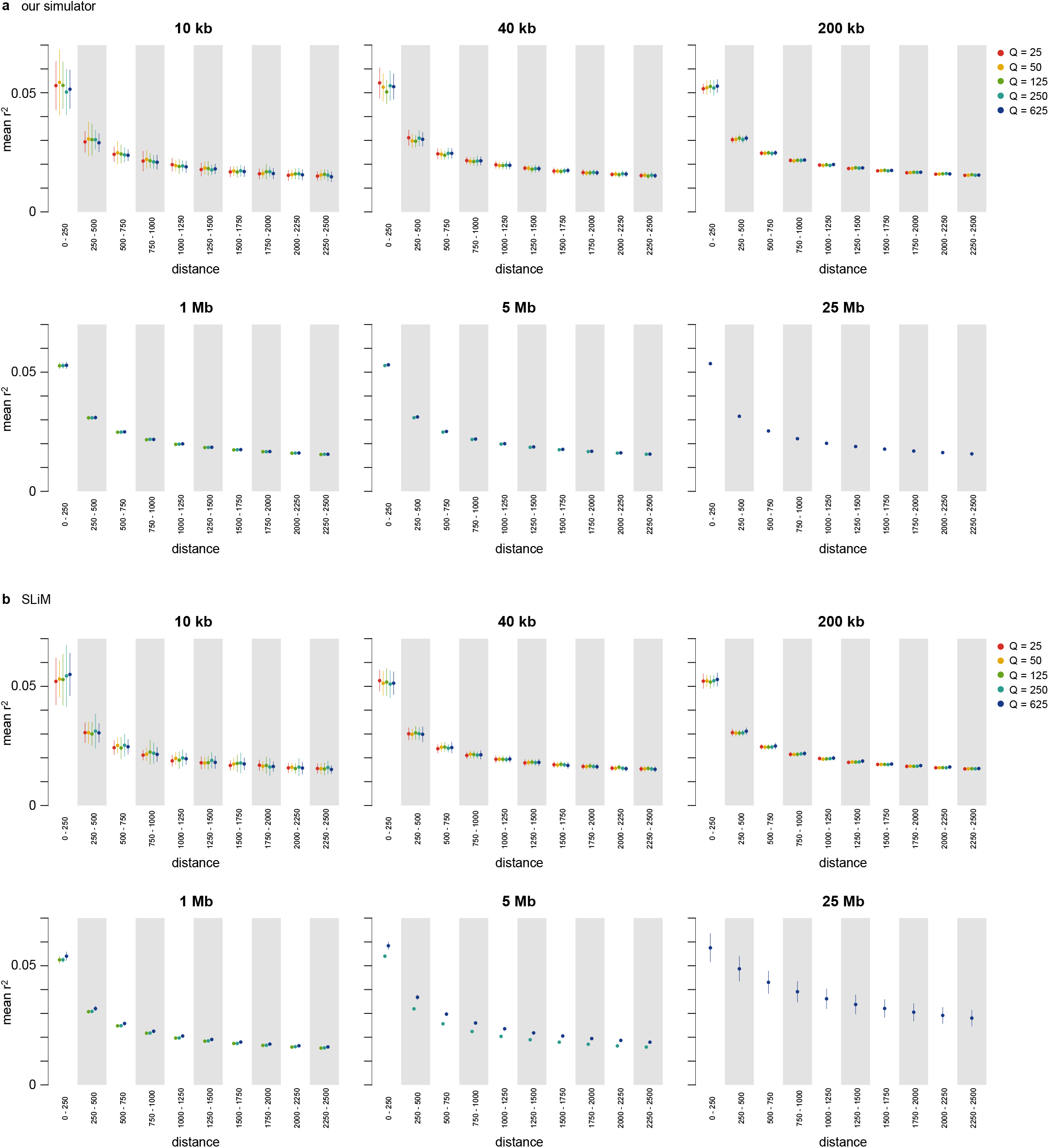
Linkage disequilibrium under purifying and positive selection. Mean squared linkage disequilibrium (*r*^2^) is shown for different genomic region lengths and rescaling factors *Q*. Values were calculated in non-overlapping 250-bp bins. Points and error bars indicate the mean and standard deviation across 50 simulation replicates. Panels (a) and (b) show results from our simulator and SLiM, respectively.

**Figure S10.**
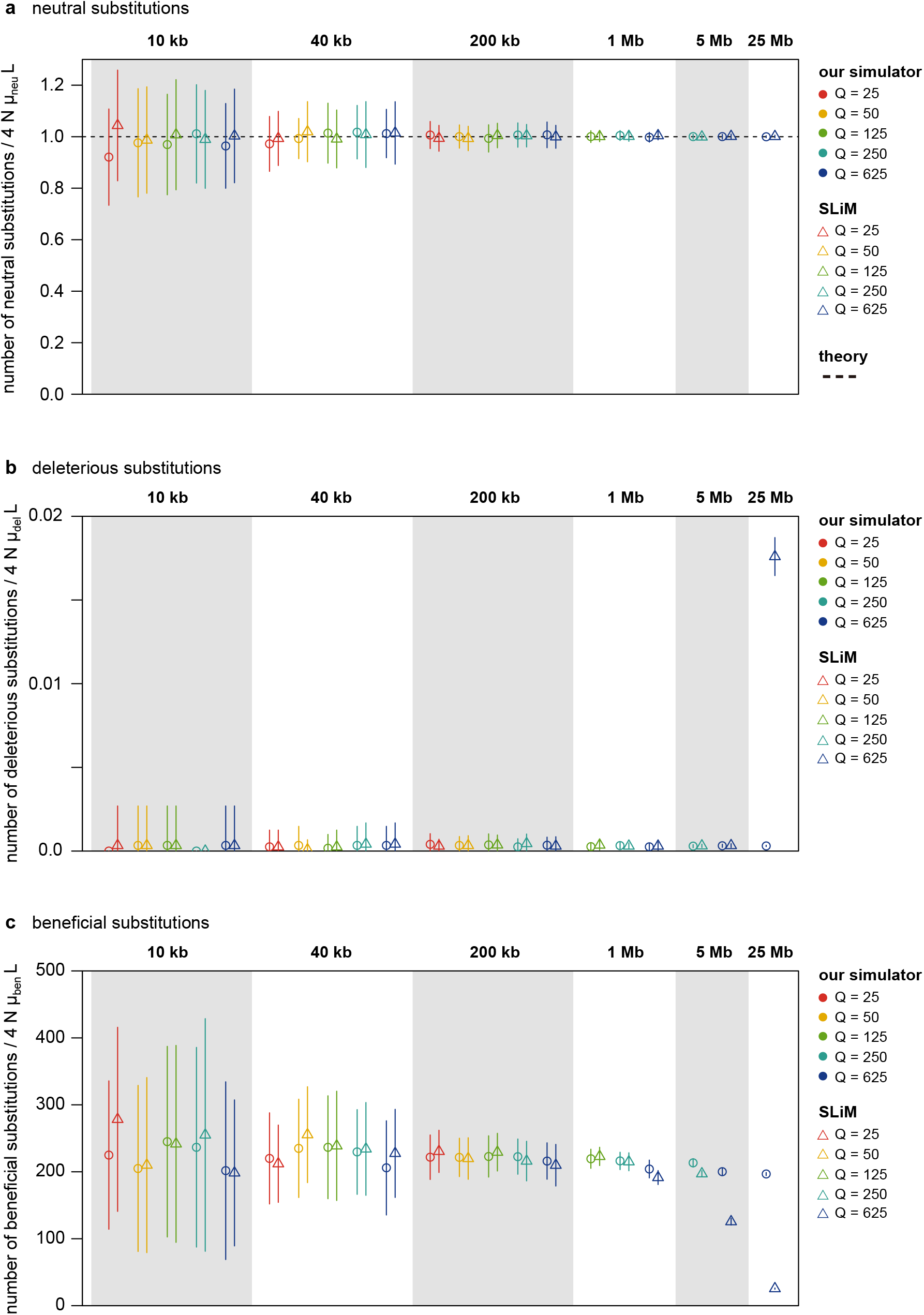
Substitution counts under purifying and positive selection. The numbers of neutral, deleterious, and beneficial substitutions fixed during the last 4*N*_rescaled_ generations are shown for different genomic region lengths and rescaling factors *Q*. Counts are normalized by the corresponding neutral expectations, 4*Nµ*_neu_ *L* for neutral substitutions, 4*Nµ*_del_ *L* for deleterious substitutions, and 4*Nµ*_ben_ *L* for beneficial substitutions. Points and error bars indicate the mean and standard deviation across 50 simulation replicates. Panel (a) shows neutral substitutions, panel (b) deleterious substitutions and panel (c) beneficial substitutions; in each panel, circles and triangles represent our simulator and SLiM, respectively.

**Figure S11.**
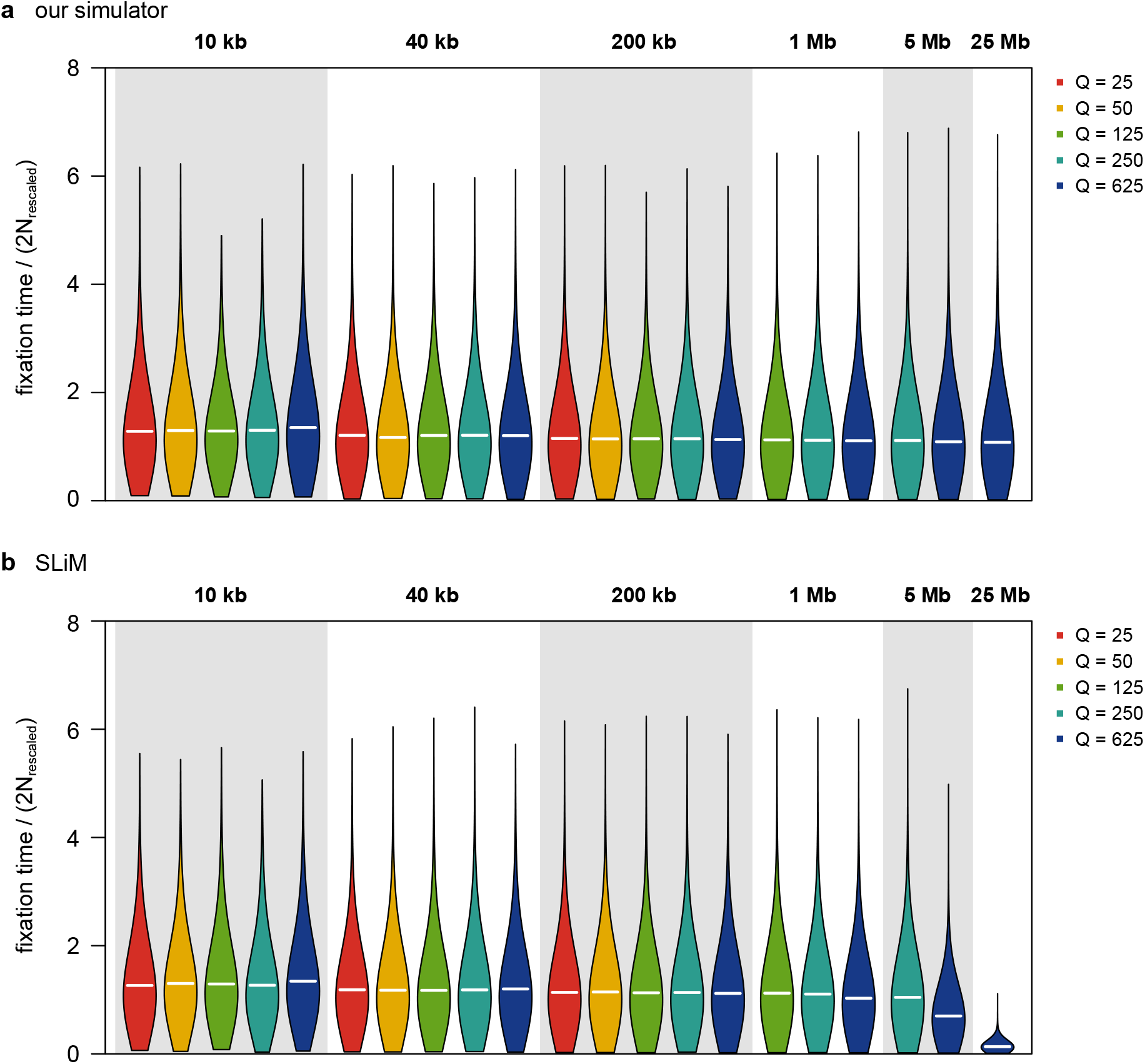
Fixation time of neutral substitutions under purifying and positive selection. Distributions of fixation times for neutral mutations fixed during the last 4*N*_rescaled_ generations are shown for different genomic region lengths and rescaling factors *Q*. Fixation times are normalized by 2*N*_rescaled_. Horizontal white lines indicate the mean fixation time. Panels (a) and (b) show results from our simulator and SLiM, respectively.

**Figure S12.**
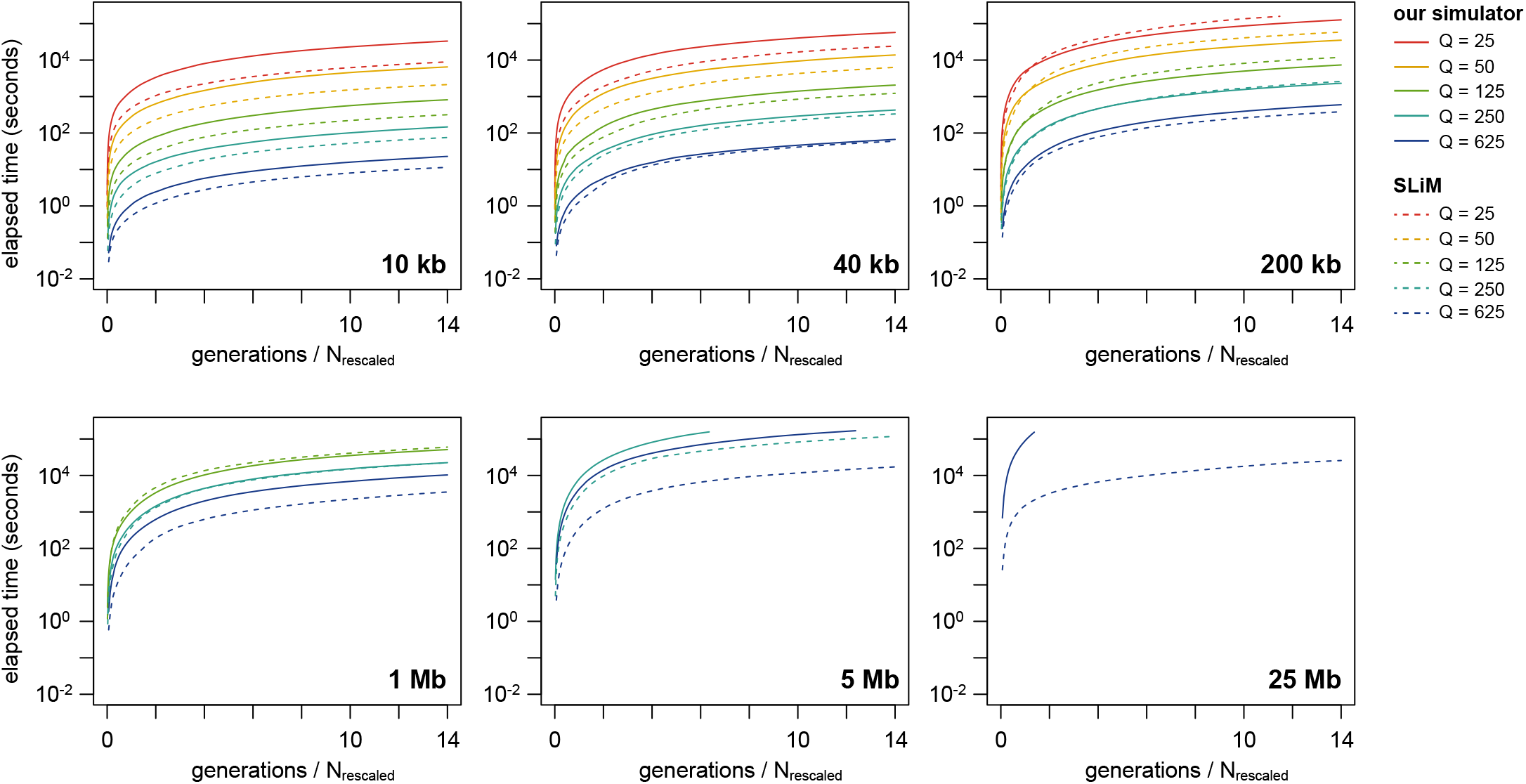
Comparison of computational time. Elapsed time is plotted as a function of the number of generation scaled by *N*_rescaled_. Median value across 10 replicates is plotted. Simulations were run at the computer cluster SHIROKANE at the Human Genome Center (the Univ. of Tokyo) using a single CPU core of Shirokane 8 (mjob).

## Footnotes

1 The parameter value reported in Marsh *et al*. (2026) contains a typographical error: it should be *f*_pos_ = 0.0002 rather than *f*_pos_ = 0.002 (J. Marsh, personal communication)

